# Atypical immunity induced by extracellular water in plants

**DOI:** 10.64898/2026.08.25.745410

**Authors:** Faye Gaudreault-Lafleur, Charles Roussin-Léveillée, Sabrina Gauthier, Gaële Lajeunesse, Alexis Roy, Isabelle Laforest-Lapointe, Peter Moffett

## Abstract

Microbial pathogens require nutrients and water to support their growth and proliferation. Pathogen-mediated resource acquisition is orchestrated by the secretion of virulence factors that have evolved a diversity of forms (from effector proteins to small toxins), but which have converged in function (resource acquisition). A key observable virulence mechanism employed by a large number of microbial pathogens to cause disease is to induce a water-rich niche in the extracellular space (i.e. the apoplast) of their host, known as ‘water-soaked lesions’. Given the ubiquity of this virulence mechanism, we asked whether plants respond to pathogen-induced extracellular water to activate immune programs. We find that inducing a water-soaked apoplast induces an atypical transcriptional response, yet results in a functional immune response that restricts pathogen growth as effectively as canonical pattern-triggered immunity. Genetic and functional analyses reveal a role for the immune phytohormone salicylic acid (SA) in mediating apoplastic water-induced immunity (AWII). These results indicate that plants engage in an immune-priming program in response to apoplastic water accumulation and suggest that the pathogenic niches induced during infection may be perceived as danger signals.

## Introduction

The threat of microbial pathogens to food security is increasing globally as a direct consequence of climate change (Singh et *al*., 2023). Among the key abiotic factors projected to increase in severity throughout the next decades are more intense rainfall following periods of drought (Guccione et *al*., 2026). High humidity during rainfall periods is associated with the onset of a large number of plant diseases, mostly of bacterial, fungal and oomcyte origin (Roussin-Léveillée et al., 2024). A major element in disease onset is the appearance of ‘water-soaked lesions’ in the extracellular space of aerial tissues under high humidity, which is essential for disease progression and has been proposed to be a key virulence mechanism employed by many pathogens (Xin *et al*., 2016).

Microbial pathogens produce effector molecules, including proteins and toxins, to interfere with the plant immune system and to induce water/nutrient-rich niches in the apoplast (Wang et al., 2022; Roussin-Léveillée *et al*., 2024). This effector-driven extracellular niche (EDEN) can be induced through a variety of means: transcriptional manipulation of host susceptibility genes, production of water/nutrient channels or manipulation of plant physiological processes (Roussin-Léveillée *et al*., 2024). In the case of the model bacterial pathogen *Pseudomonas syringae pv. tomato* DC3000 (*Pst*), water-soaking is induced by manipulation of host abscisic acid (ABA) biosynthesis and signaling, leading to stomatal closure and therefore, accumulation of apoplastic water (Hu *et al*., 2022; Roussin-Léveillée *et al*., 2022). These water-soaked lesions are induced by *Pst* through the secretion of two type III effector proteins, HopM1 and AvrE1, which modulate ABA-related pathways (Hu *et al*., 2022; Roussin-Léveillée *et al*., 2022). Manipulation of ABA signaling by microbial effectors has been reported in multiple pathosystems, converging on immune inhibition and water-soaking functions (Mine *et al*., 2017; Peng *et al*., 2019; You *et al*., 2023; Herold *et al*., 2025; Joshi *et al*., 2026).

To prevent infections, plants have evolved to sense pattern-associated molecular patterns (PAMPs) through large repertoires of plasma-membrane localized pattern-recognition receptors (PRRs) in the apoplast (Ngou *et al*., 2022). Upon PAMP perception by PRRs, plants activate signalling cascades referred to as pattern-triggered immunity (PTI), which contributes to disease resistance (DeFalco *et al*., 2021). PRRs also respond to normally sequestered, free-flowing compounds, such as cellotriose or other types of sugars, in the apoplast associated with cell wall damage induced by pathogens (Martín-Dacal *et al*., 2023; Jiménez-Sandoval *et al*., 2025). In this scenario, damage-associated molecular patterns (DAMPs) induce a response known as DAMP-triggered immunity (DTI) (Harris *et al*., 2024). Cell surface PRRs function through association with co-receptors at the plasma-membrane, such as BRASSINOSTEROID-ASSOCIATED KINASE1 (BAK1) and other SERK-family types of receptor kinases (Wu *et al*., 2025). In turn, PTI/DTI activation leads to transcriptional immune reprogramming, defense hormone accumulation and cell wall/apoplastic-based defenses (DeFalco *et al*., 2021).

The withholding of apoplastic water, or water immunity, appears to be a key element of plant defenses (Jian *et al*., 2023; Kemppinen *et al*., 2025). We have previously reported that plants can suppress pathogen-induced water-soaking when the salicylic acid (SA) branch of plant immunity is activated either by exogenous application or by photoperiod stress (Lajeunesse *et al*., 2023). Suppression of water-soaking also appears to be a key defense mechanism in plants undergoing effector-triggered immunity (ETI), an immune reaction triggered by intracellular recognition of pathogenic effectors by NLR proteins (Xin et al., 2016; Schenstnyi *et al*., 2022).

Given the importance of water for pathogen growth and the ubiquity of water-soaking in plant-microbe interactions, we hypothesized that extracellular water may act as a danger signal in plants. In this report, we assess the *Arabidopsis thaliana* transcriptional and physiological responses to extracellular water, which we show to mimic pathogen-induced water-soaking. Our results demonstrate that extracellular water triggers an atypical transcriptional immune response in Arabidopsis, which leads to effective SA-dependent disease resistance. This response is similar in magnitude to the effects of typical PTI responses and provides an effective protection against pathogen infection.

## Results

### Apoplastic water rewires the Arabidopsis transcriptional landscape

To evaluate whether apoplastic water alone induces a response in plants, we performed RNA-sequencing on *Arabidopsis thaliana* leaves infiltrated with a control solution, *Pst* WT or *Pst hrcC*^−^ (type-III secretion system mutant defective in water-soaking induction). To distinguish between PTI, water-soaking and other *Pst*-mediated effects on plant transcriptomes, we either maintained the apoplast flooded (F) with the infiltrates (water, *Pst* WT or *hrcC^−^*) or allowed it to return to the pre-infiltration state (non-flooded; NF) (Fig. 1A). Interestingly, we find that the transcriptome of Arabidopsis leaves diverged when the apoplast is maintained in an F state compared to an NF state only with water or *Pst hrcC*^−^ but not with *Pst* WT (Fig. 1B-D). This could be explained by the fact that *Pst* WT induces water-soaking under high humidity and therefore, the apoplast of NF state plant infected with *Pst* WT was naturally flooded by these lesions. Importantly, this observation indicates that artificially maintaining the apoplast in a F state with *Pst* WT does not lead to further unexpected transcriptional responses that do not occur in plants undergoing water-soaking lesions. Gene ontology (GO) analysis of genes overexpressed in the F state of control (water) or *Pst hrcC*^−^ infiltrated compared to their NF state counterpart highlights an overrepresentation of GO terms associated with immune responses and biotic stress sensing (Fig. 1E,F).

**Figure 1:**
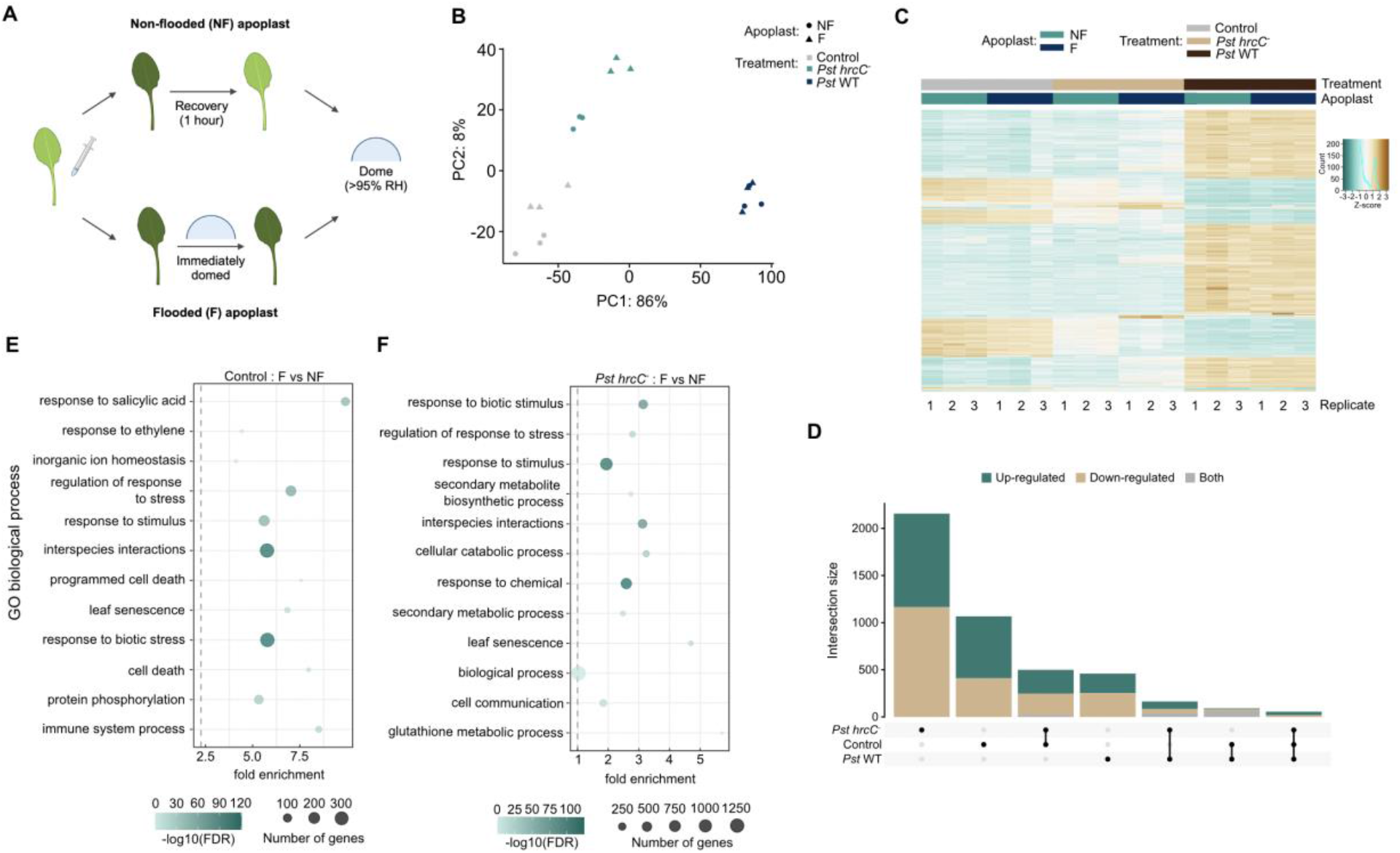
Apoplastic water alters the Arabidopsis transcriptional landscape. (**A**) Schematic of the experimental setup to evaluate the impact of apoplastic water flooding on Arabidopsis leaf transcriptional program. Leaves were either maintained in a flooded (F) state by covering with a plastic dome immediately after infiltration to maintain high relative humidity and prevent evapotranspiration, or plants were left undomed, allowing excess moisture to evaporate (non-flooded; NF). In both cases, plants were maintained under high humdity for 24 hours prior to tissue harvesting. Created with Biorender. (**B**) Principal component analysis showing expression changes across Arabidopsis leaves in response to apoplast flooding under control and infected conditions. Four-week-old Arabidopsis leaves were infiltrated with either MgCl_2_ 10 mM (control), *Pst* WT or *hrcC^−^* (1×10^5^ CFU/ml) and maintained either in a F or NF state as mentionned in A. Leaves were harvested 24 hours post treatment and RNA extracted before being subjected to RNA-seq. (**C**) Z-score heatmap displaying how gene expression is affected across apoplastic conditions and treatments and between biological replicates from RNA-seq results obtained in A. Rows and columns are grouped using hierarchical clustering to reveal co-expressed gene modules. Colors represent row-standardized Z-scores. (**D**) Intersection of extracellular water-responsive DEGs across bacterial infection contexts. UpSet plot showing the overlap of differentially expressed genes (DEGs) identified in three independent within-group F vs NF states. (**E-F**) Gene ontologies enriched in the of the F state compared to the NF state in Arabidopsis leaves infiltrated with MgCl_2_ 10 mM (control) (E) or *Pst hrcC^−^*(1×10^5^ CFU/ml) (F) as in B.

We next evaluated the gene-regulatory network structure of the response to extracellular water presence in uninfected plants. Analysis of the top transcription factors (TFs) upregulated by extracellular water revealed a diversity of transcription factor families, including members from the WRKY, ERF and NAC families (Fig. S1A). We found a clear enrichment in WRKY-binding motifs in DEGs associated with extracellular water responsiveness (Fig. S1B). To prioritize candidate master regulators of the extracellular water transcriptional response, we used a regulatory potential score that integrates the magnitude of TF induction, the statistical enrichment of its binding motif in the promoters of extracellular water-responsive genes, and the effect size of that enrichment. TFs were ranked by this composite score, with WRKY TFs clearly dominating the top candidates (Fig. S1C). Considering the impact of WRKY TFs in early immune signaling, it is not suprising to see that WRKY family members dominate the predicted regulatory hierarchy of the extracellular-water response.

### Atypical, but functional, immune response elicited by apoplastic water

As GO terms associated with biotic stress responses and immunity were significantly enriched in our flooded apoplast transcriptomes (both control or in the presence of *Pst hrcC*^−^), we sought to evaluate whether this response was enough to confer disease resistance after pathogen challenge. To evaluate this, we designed an experiment in which we maintained the apoplast flooded (F) for 20 hours and allowed it to return to the pre-infiltration state for 4 hours prior to being challenged with a bacterial infection. Previously untouched plants or plants infiltrated with water 20 hours prior to infection but were allowed to immediately return to a pre-infiltration state (NF) served as negative controls. As a positive control, plants were pre-treated with the immunogenic flagellin epitope flg22, which is known to confer protection (Zipfel *et al*., 2004). As expected, three days after inoculation, untouched and NF leaves showed strong disease symptoms (Fig. 2A). However, disease symptoms were largely absent in leaves that were either pretreated with flg22 or maintained in a F state for 20 hours prior to inoculation (Fig. 2A). Consistent with differences in disease symptoms, both flg22 and F pre-treated leaves showed similar significant reductions in *Pst* WT growth compared to untouched and NF leaves (Fig. 2B). We refer to this phenomenon as apoplastic water-induced immunity (AWII) hereafter. We next investigated the temporal dynamics of AWII establishment. We found that shorter F pre-treatment durations (2, 4 or 6 hours of F state) were insufficient to induce AWII compared to the 20 h F pre-treatment, as indicated by the ability of *Pst* to induce water-soaking symptoms (Fig. 2C) or restriction of bacterial growth (Fig. 2D). This suggests that plants do not simply activate AWII upon a temporary hydration of the apoplast, but rather following an abnormal, longer hydration of the extracellular space, which is often associated with the presence of a pathogen.

**Figure 2:**
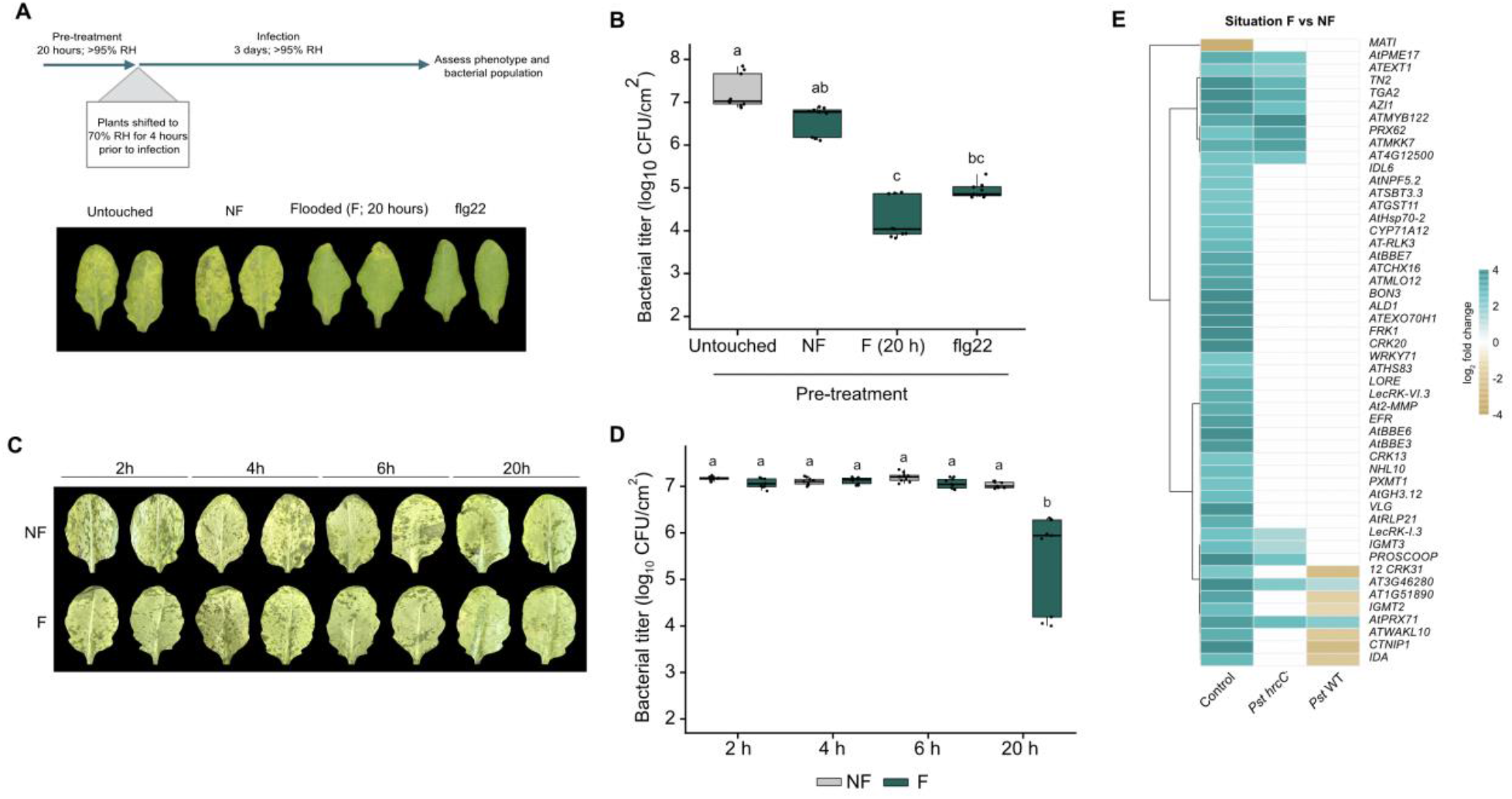
Apoplastic water pre-treatment confers protection against *Pst*-mediated disease. (**A**) Diagram of the experimental setup to evaluate the impact of apoplastic water flooding on plant disease progression and photos of Arabidopsis leaves at 3 days post-inoculation (dpi) with *Pst* WT (1 × 10^5^ CFU/ml). Plants were either untouched 24 hours prior to *Pst* infection (untouched), or infiltrated with 10 mM MgCl_2_ and were either allowed to return to a non-flooded state (NF) or water was maintained in leaves for 20 hours prior to returning to a pre-infiltration state (flooded; F), or treated with the immunogenic peptide 1 μM flg22. In all cases, plants were maintained under high humdity for the first 20 hours and switched to low humidity for the last 4 hours prior to infection. (**B**) Bacterial titer in plants infected as in (A) at 3 dpi. (**C**) Disease protection dynamics over time in plants that were maintained in a F state. Leaves were treated as in A, but with a F state for 2, 4, 6 or 20 hours prior to being challenged *Pst* WT (1 × 10^5^ CFU/ml). Photos were taken 24 hours post inoculation (hpi) (**D**) Bacterial titer in plants infected as in (C) at 3 dpi. (**E**) Heatmap representing expression differences of genes associated with pattern-triggered immunity in response to artificial apoplast flooding across different scenarios. Different letters indicate statistical significance (p<0.05) obtained after performing a Kruskal-Wallis followed by a Dunn’s post-hoc (BH correction) (2B, 2D).

Since AWII is as effective as flg22 in mediating disease resistance against *Pst* WT, we evaluated whether AWII activates a PTI-like transcriptional response. We evaluated the expression pattern of PTI marker genes in Arabidopsis leaves that were kept in a F vs NF state in the absence of bacteria (control), or in the presence of either *Pst hrcC*^−^ or *Pst* WT. Interestingly, the greatest differential in upregulation of PTI-related genes was observed in the control F condition compared to control NF (Fig. 2E). In leaves inoculated with WT or mutant *Pst*, there was little differences between F and NF conditions likely because PTI-related genes are induced in the NF condition upon perception of *Pst* and are therefore not significantly affected by apoplastic water retention.

We next compared the transcriptome of Arabidopsis plants in a F vs NF state at 20 hours post water infiltration (no bacteria) to an extensive transcriptional study of the PTI/DTI response in plants (Bjornson *et al*., 2021). While PTI induced by the immunogenic elicitors flg22, elf18, nlp20, ch8 and LPS or DTI elicitors Pep1 and OGs share relatively conserved transcriptional signatures in Arabidopsis seedlings, only a small subset of genes regulated during AWII was found to overlap with the PTI/DTI transcriptional signature (Fig. S2). This suggests that AWII results in a transcriptional response that differs significantly from responses associated with plasma-membrane perception of immune eliciting compounds. Nevertheless, water accumulation in the apoplast does appear to act as an atypical immune cue that promotes transcriptional response that overlaps with PTI responses and ultimately results in a similar outcome.

### Apoplastic water-induced immunity is independent of hypoxia, mechanosensing and senescence signalling

Upon comparison of the AWII transcriptional response to PTI/DTI responses, we were able to identify AWII-specific gene clusters that are not regulated by either PTI or DTI. GO term enrichment on this specific subset revealed gene categories associated with energy and translation being significantly downregulated (Fig. 3A). This could be caused by a hypoxic stress that limits ATP production, forcing plants to slow down high-energy processes such as transcription and ribosome assembly. Flooding the apoplast alone was sufficient to drive a clear transcriptional response resembling that seen in a hypoxia response driven by the ERF-VII factor (Licausi *et al*., 2011). Interestingly, all treatments showed a certain overlap with this response (Fig. 3B). Indeed, *Pst hrcC*^−^ in the flooded state showed a response similar to control flooded, as might be expected. At the same time, we observed a clear induction of a hypoxia-related response in response to *Pst hrcC*^−^ non-flooded conditions, suggesting that the PTI response induced by this attenuated strain overlaps with the hypoxia response (Figs. 3B, S3). Likewise, *Pst* WT induced hypoxia-responsive genes to an even greater extent (Fig. 3B), probably due to water-soaking and a reduction in plant transpiration. At the same time, certain hypoxia-related genes were not induced, or were down-regulated by *Pst* WT infection, possibly due to the defense-suppressive effects of this pathogen (Figs. 3B, S3).

**Figure 3:**
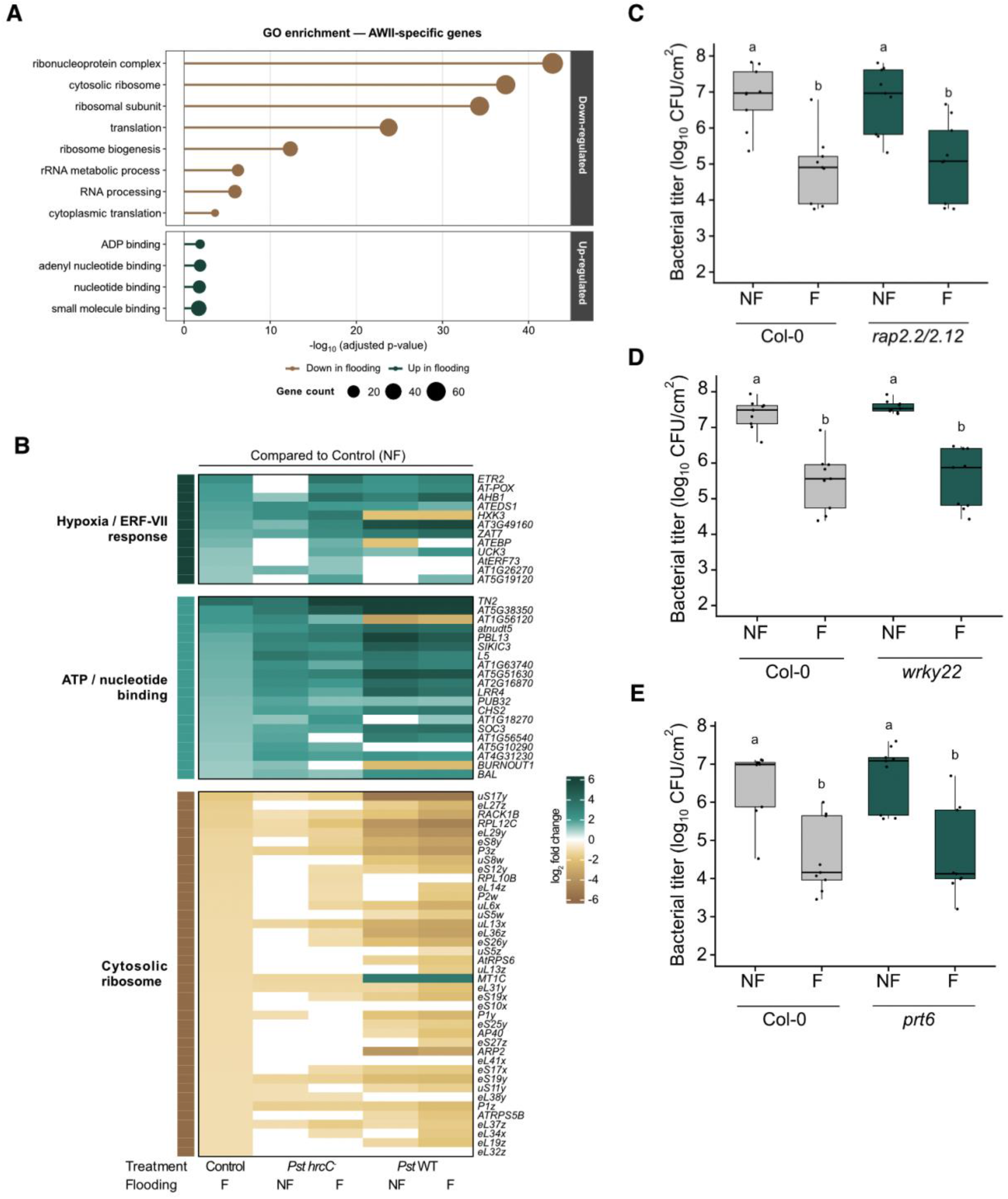
Apoplastic water-induced immunity is not triggered by hypoxia. (**A**) Gene Ontology enrichment of AWII flooding-specific genes. Lollipop plot showing enriched Gene Ontology (GO) terms for the 575 genes regulated by apoplast flooding in the AWII transcriptome (control; NF vs F) that were absent from the DEG sets of all seven P/DAMP treatments in the Bjornson *et al*., (2021) dataset. Dot size is proportional to the number of genes from the test set annotated to each term. (**B**) Flooding transcriptional response of AWII-specific genes across bacterial infection contexts. Heatmap displaying log₂ fold change values for 74 differentially expressed genes (DEGs) identified as AWII-flooding-specific, defined as genes significantly regulated by the F state in Arabidopsis WT apoplast that were absent from the filtered DEG sets of all seven P/DAMP treatments from the Bjornson *et al*., 2021 dataset (Figure S2). (**C-E**) Hypoxia is not involved in the AWII effective response. Bacterial titer at 3 dpi in Arabidopsis hypoxia-related mutants *rap2.2/rap2.12* (C), *wrky22* (D) and *prt6* (E) that were subjected to AWII (F state for 20 hours) or NF state as exemplified in 2A, prior to being challenged with *Pst* WT (1 × 10^5^ CFU/ml). Different letters indicate statistical significance (p<0.05) obtained after performing a one-way ANOVA followed by a Tukey HSD post-hoc (3C), Welch’s ANOVA followed by a Games-Howell post-hoc (3D) and a Kruskal-Wallis followed by a Dunn’s post-hoc (BH correction) (3E).

To evaluate whether AWII was caused by the impact of hypoxia on plant cells, we tested if AWII could be induced in plants defective in hypoxia responses. Physiological hypoxia is regulated by the Group VII Ethylene-Responsive Transcription Factor (ERF-VII) family (Licausi *et al*., 2011). RAP2.2 and RAP2.12 are two ERF-VII transcription factors which act redundantly and are required for inducing hypoxia-induced transcriptional responses (Licausi *et al*., 2011). We find that apoplast flooding was still able to induce a protective effect against bacteria in an Arabidopsis *rap2.2/rap2.12* mutant (Fig. 3C). This suggests that the initial transcriptional hypoxia response is not necessary for AWII execution. Hypoxia can also be caused by environmental conditions, such as submergence/flooding. Environmental hypoxia caused by submergence has been shown to confer immunity and this response requires the transcription factor *WRKY22* (Hsu *et al*., 2013). We found that Arabidopsis *wrky22* mutants still induces AWII to levels similar to WT plants (Fig. 3D), indicating that leaves do not engage a submergence-induced immune program under such conditions. To further rule out the implication of hypoxia in the AWII response, we used an Arabidopsis mutant for the *PROTEOLYSIS 6* (*PRT6*) gene, which is a central negative regulator of the hypoxia response and *prt6* mutants induce a hypoxia response even under non-hypoxic conditions (Gibbs *et al*., 2011). However, *prt6* mutants show *Pst* growth levels similar to WT Arabidopsis, indicating that a constitutive hypoxia response does not induce an AWII-like protective response against bacterial infection (Fig. 3E). Likewise, the flooding pretreatment in the *prt6* mutant induced a protective effect similar to WT Arabidopsis, indicating that this mutant is still competent for AWII (Fig. 3E). Together, results indicate that hypoxia is unlikely to be the central molecular mechanism behind AWII activation.

Extracellular water could potentially affect other plant processes, such as membrane mechanics, liberate immune-eliciting cell wall components and activate early senescence. The Arabidopsis genes *MID1-COMPLEMENTING ACTIVITY 1* (*MCA1*), *HAESA-LIKE 3* (*HSL3*) and ALTERED HYDROTROPIC RESPONSE 1 (*AHR1*) are involved in mechano-transduction, phytocytokine recognition in water-stress responses and sensing of water potential changes, respectively (Nakagawa *et al*., 2007; Saucedo *et al*., 2012; Rhodes *et al*., 2022;). We found that AWII could still elicit a strong protection in Arabidopsis single mutant plants for each of these genes (Fig. S4A-C), indicating that they are not required for AWII responses. Submergence-associated oxygen depletion is known to trigger early senescence and the transcription factor *ORESARA1* (*ORE1*) is required for this process (Rankenberg *et al*., 2024). Similarly to what was observed in other Arabidopsis mutants, we found that AWII was still fully functional in preventing bacterial proliferation in the Arabidopsis *ore1* mutant (Fig. S4D). Taken together, our results indicate that certain plausible mechanisms such as hypoxia, mechano-sensation and senescence, that could be activated by the presence of extracellular water are not associated with the establishment of AWII.

### Apoplastic water-induced immunity is salicylic acid-dependent

The defense hormone salicylic acid (SA) is required for multiple defense-related responses (Spoel and Dong, 2024) including the blocking of water-soaking (Lajeunesse *et al*., 2023). We therefore asked whether AWII could be a SA-driven priming mechanism. Analysis of SA-associated gene expression during AWII showed that the SA biosynthetic genes (*ICS1/SID2*) and biosynthesis-regulating genes (*CBP60g* and *SARD1*) are up-regulated during AWII, albeit to a lesser degree than treatments including *Pst* WT or *hrcC^−^*, but that downstream SA signaling genes (*PR* genes) were not up-regulated (Fig. 4A). Ethylene also operates at the intersection between hypoxia and immunity signaling in plants (Huang *et al*., 2016; Hartman *et al*., 2021). Evaluation of ethylene-responsive genes during AWII revealed very little regulation of the ethylene pathway by the presence of extracellular water, suggesting that SA is more likely than ethylene to be involved in mediating AWII (Fig. 4A). We next evaluated SA accumulation during AWII in Arabidopsis leaves. We found that leaves undergoing AWII (F state) accumulated slightly more SA than plants kept in a NF state, whereas treatment with flg22 resulted in a marked increase in the accumulation of SA (Fig. 4B). Interestingly, when leaves were infiltrated with flg22 and kept in a F state, this led to a nearly two-fold increase in SA accumulation compared to flg22 treatment without flooding (Fig. 4B), suggesting a possible additive effect of the two treatments.

**Figure 4:**
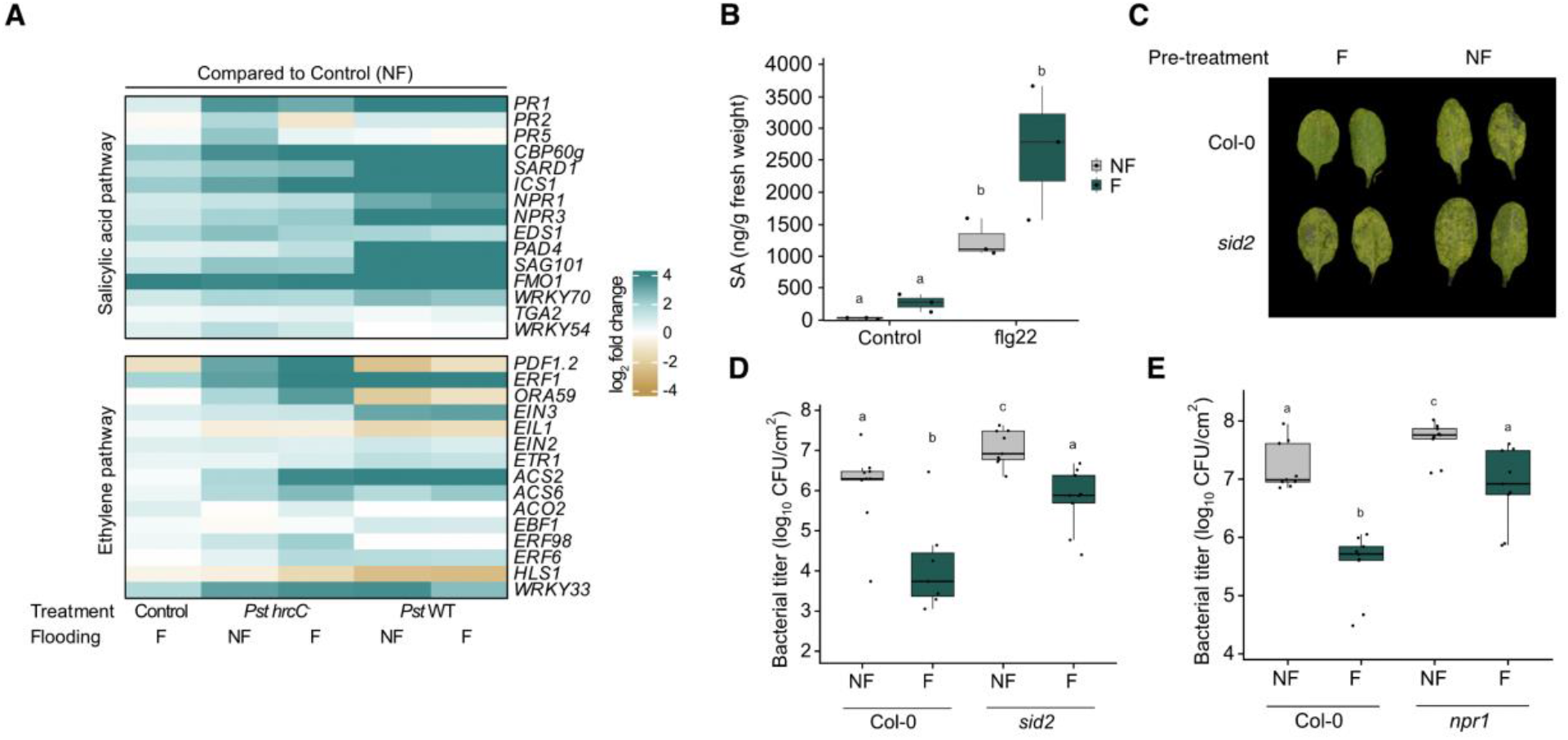
Apoplastic water-induced immunity is salicylic acid-dependent. (**A**) Heatmap representing expression differences of genes associated with the salicylic acid (SA) and ethylene defense hormone pathways. Log_2_ fold change in expression differences between Control F, *Pst hrcC^−^* and *Pst* WT in the NF and F states compared to Control NF of each individual treatment from the RNA-seq presented in 1B. (**B**) Apoplast flooding leads to an increase in SA accumulation in the presence or absence of flg22. Quantification of SA at 20 hours post treatment by UPLC-MS from Arabidopsis leaves that were either infiltrated with 10 mM MgCl_2_ or 1 μM flg22 that were maintained in NF or F state. Different letters indicate statistical significance (p<0.05) after performing a one-way ANOVA followed by Tukey’s HSD post-hoc. (**C**) Deficient AWII response in the Arabidopsis *sid2* mutant. Four-week-old Arabidopsis WT or *sid2* leaves were infiltrated with 10 mM MgCl_2_ and either maintained in a NF or F state for the first 20 hours prior to being removed from high humidity conditions for 4 hours and challenged with *Pst* WT (1 × 10^5^ CFU/ml). Photos were taken at 3 dpi. (**D-E**) Bacterial titer in plants treated as in (C) in the Arabidopsis *sid2* (D) or *npr1* mutant plants at 3 dpi. Different letters indicate statistical significance (p<0.05) obtained after performing a one-way ANOVA followed by a Tukey HSD post hoc test (4B) or a Kruskal-Wallis followed by a pairwise Wilcoxon with BH correction (4D, 4E).

To evaluate the role of SA in mediating the AWII response, we tested AWII execution in the Arabidopsis SA biosynthesis mutant *sid2-2* (*sid2*) and the SA receptor/transductor mutant *npr1-1* (*npr1*) (Wildermuth *et al*., 2001; Wu *et al*., 2012). We found that in contrast to Arabidopsis WT leaves, *sid2* mutant leaves pretreated with flooding showed strong disease symptoms three days after bacterial inoculation (Fig. 4C). *Pst* WT bacterial titers in *sid2* plants having undergone AWII showed slightly lower bacterial growth in F conditions compared to NF. However, the differential was much less than that seen in WT plants and *Pst* grew to levels similar to those observed in WT plants that were not protected by AWII (Fig. 4D). Similar effects were seen on bacterial growth in *npr1* mutants (Fig. 4E). Together, these results demonstrate that apoplastic water-induced immunity requires functional SA biosynthesis and signaling transduction, suggesting that AWII likely shares certain mechanistic elements with other types of defense responses.

## Discussion

Pathogen-induced water-soaking lesions are common symptoms associated with the onset of pathogenesis in plants and have been described as serving the pathogen, providing the moisture it needs to proliferate (Aung *et al*., 2018; Roussin-Léveillée *et al*., 2024). In this study, we provide evidence that extracellular water, in addition to being a key component of virulence, can also activate a previously unreported type of immune response. We show that AWII activates immune-associated transcriptional programs as well as functional priming of protection against subsequent infections. We find that this specific type of priming is SA-dependent, despite SA not accumulating significantly in the absence of a pathogen. Our findings raise an interesting question for understanding pathogenic niches: can plants detect extracellular modifications associated with pathogenesis and mount functional immune responses to them?

Extracellular water does not appear to be perceived as a classical danger signal through known P/DAMP-recognition receptors as its transcriptome only partially overlaps with canonical PTI and DTI signatures (Bjornson *et al*., 2021). While this could be attributed to comparing transcriptomes from different studies at different time points, PAMPs are known to induce major transcriptional changes, which do not seem to be present in the AWII transcriptome. Notably, most PAMPs engage clear SA-responsive regulatory nodes and increase SA accumulation in leaves. This is not the case for AWII, although the latter does require SA. Thus, while diverging in transcriptional outputs, AWII and PTI both lead to disease resistance, suggesting that functional priming requires only modest transcriptional changes to be effective.

How extracellular water activates immune priming is still unclear. It is notable that the effects of AWII require an extended length of time of flooding to induce a protective ffect (Fig. 2C,D) suggesting either a requirement for an extended period of sensing and/or signaling. This also begs the question of whether plants undergoing AWII respond to water via a receptor-like protein or whether AWII is a response to physiological changes. The AWII-induced transcriptional signature could be suggestive of hypoxia as being a key element of the AWII response. However, hypoxia has been reported to act as a negative regulator of immune responses in Arabidopsis (Mooney *et al*., 2024) and AWII was not altered in hypoxia response mutants (Fig. 3). However, not all types of hypoxia responses have been shown to negatively regulate plant immunity. Indeed, whole plant submergence has been shown to confer *WRKY22*-dependent resistance against *Pseudomonas syringae* (Hsu *et al*., 2013). Our results indicate that AWII does not require *WRKY22* (Fig. 3D), suggesting differing mechanisms. Submergence has also been shown to deactivate jasmonic acid-dependent, wound-induced defense against herbivores (Lee *et al*., 2020). suggesting that the relationship between hypoxia and immunity depends on which hormonal branch and which type of biotic threat is being considered (García *et al*., 2024). At the same time, it is not clear to what degree the transcriptional response seen in AWII is a direct reflection of hypoxia as we have not established the extent of gas exchange or the degree of hypoxia experienced by cells undergoing AWII. Furthermore, it is noteworthy that some hypoxia response genes are also upregulated in non-flooded conditions, in the presence of a non-virulent pathogen (Fig. 3B). This suggests that some of the hypoxia transcriptional signature may be due to overlap between hypoxia and defense responses. Ultimately however, the hypoxia response does not appear to be sufficient or necessary for the SA-dependent AWII response and may be induced independent of and/or as a consequence of defense responses.

Extracellular water could induce a physical stress on plant membranes which could, in turn, potentially activate transcriptional responses. Arabidopsis can perceive and signal membrane-associated stress and depolarization through MCA1 and MCA2, two plasma-membrane proteins required for mechanically induced Ca^2+^ influx (Gilliard *et al*., 2026). In this study, we found that AWII-mediated priming was still intact in the Arabidopsis *mca1* mutant, which is a significant mechanosensitive mutant in its own right (Nakagawa *et al*., 2007). Although we cannot rule out redundancy with other mechanosensitive channels, such as MCA2, it is notable that MCA1/MCA2 confer resistance to a necrotrophic pathogen (Gilliard, *et al*., 2026), while turgor pressure and other types of membrane stress induce jasmonic acid signalling (Hamel *et al*., 2024; Mielke *et al*., 2021). As such, it may be less likely that AWII is mediated by similar mechanisms.

Plant immunity is often described in terms of changes in gene expression, but it is equally important to consider changes in the extracellular space, where most bacterial, fungal and oomycete pathogens colonize, which may make for a less hospitable environment (Armer and van der Hoorn, 2026). Immune activation can lead to inhibition of water-soaking (Xin *et al*., 2016; Lajeunesse *et al*., 2023) and a major element of SA-dependent defence mechanisms involves keeping stomata open to prevent water transpiration (Lajeunesse *et al*., 2023). Indeed, a recent study has reported that water immunity, defined as the plant’s capacity to prevent water-soaking, plays a more prominent role than stomatal immunity in resistance to bacterial infection (Kemppinen *et al*., 2025).

The description of apoplastic water-induced immunity above raises questions regarding its potential role in plant-microbe interaction outcomes. Our finding that apoplastic water primes defenses via the SA pathway, further highlights the foundational role of this defense phytohormone as an anti-EDEN agent. Future work will be required to understand how extracellular water is perceived and to what extent it contributes to disease resistance. As with investigation of PTI signaling by canonical P/DAMPs, AWII is likely masked in the context of a compatible infection due to the defense-suppressing effects of pathogen effectors. At the same time, unlike PAMP pretreatment, the fluctuation between flooded and non-flooded apoplast may resemble realistic conditions. That is, water-soaking often occurs during more humid conditions at night, and disappear during the day (Clayton, 1936). As such, it is possible that under fluctuation environmental conditions, plants would be subjected to AWII-like conditions, which in turn may allow plants to overcome initial infections.

## Author contributions

C.R.-L. conceived the study. F.G.-L. and C.R.-L. performed all experiments, data analysis and the creation of figures equally. S.G., G.L. and A.R. contributed to some experiments and data collection. F.G.-L. and C.R.-L. wrote the first draft of this manuscript. F.G.-L., C.R.-L. and P.M. contributed to final editing of the manuscript. C.R.-L. and P.M. supervised the study. P.M. and I.L.-L. acquired research funds. All authors revised and agreed with the content written in this manuscript.

## Acknowledgments

This study was supported by a Natural Sciences and Engineering Research Council of Canada (NSERC) Discovery Grants to P.M. C.R.-L. was supported by a VoiceAge excellence fellowship and an FRQ-NT PhD scholarship.

## Materials and Methods

### Plant growth conditions

Arabidopsis Col-0 (wild-type; WT) and mutant derivatives thereof, plants were grown in PromixTM soil (PremierTech) in growth chambers with a 12 h light/dark photoperiod, with relative humidity of ∼60% at 21 °C. Light intensity was measured at 180 µmoles/m^2^/s when lights were on. Four- to five-week-old Arabidopsis plants were used for all experiments described herein, unless stated otherwise. The following mutant lines were obtained from the Arabidopsis biological resource center (Ohio State University): *hsl3* (SALK_207895C), *ore1* (CS_859988), *mca1* (SALK_206846C), *ahr1* (SALK_066191C).

### Bacterial disease assays

*Pseudomonas syringae pv. tomato* DC3000 WT and mutant strains were cultured overnight at 28 °C in Luria-Bertani (LB) media containing 50 mg/l of rifampicin. On the day of the infection, fresh LB media was inoculated with 0.5 ml of the overnight culture and bacteria were collected when OD_600_ reached between 0.8–1. Bacteria were centrifuged for 10 min at 4000 × g and the pellet resuspended in MgCl_2_ 10 mM. Bacterial density was adjusted to 0.2 (1 × 10^8^ CFU/ml) prior to further dilutions. Bacterial infections were carried out between 14:00–15:00 (zeitgeber time of 6:00–7:00).

All inoculations of Arabidopsis leaves were performed by syringe-infiltration. Infiltrated plants were all kept under ambient humidity levels for 1 h to allow water to evaporate, then domed with a plastic unit to maintain high humidity (>95% RH). Artificial flooding was induced by treating the plants in the same manner, except that plants were immediately domed after infiltration to avoid evaporation of excess liquid in the extracellular space.

Bacterial growth *in planta* was monitored by harvesting infected Arabidopsis leaves, surface sterilizing in 80% ethanol and rinsing in sterile water twice. Leaf disks were taken from three leaves from the same plant (one per leaf; total of three leaf disks) using a cork borer (6 mm in diameter) and ground in sterile 10 mM MgCl_2_. Three biological replicates were performed for each experiment. Colony-forming units (CFU) were determined by making serial dilutions (10^0^−10^−6^) and plating on LB plates containing 50 mg/l of rifampicin. Each dilution was plated in three technical replicates.

### RNA extraction and sequencing

Arabidopsis leaves were harvested and immediately flash-frozen in liquid nitrogen prior to RNA extraction. RNA was extracted from flash-frozen, ground leaf tissue followed with QIAZOL (QIAGEN) reagents followed by on-column DNase treatment (QIAGEN), according to the manufacturer’s protocol. RNA integrity was evaluated by an Agilent Bioanalyzer 2100 with the Eukaryote Total RNA Nano Series II. cDNA libraries were generated using NEBNext Ultra II Directional RNA Library Prep Kit for Illumina kit (New England Biolabs, USA) according to the manufacturer’s protocol. cDNA libraries were sequenced by RNA-seq at the Université de Sherbrooke RNomics Platform using an Illumina NextSeq 500 system. Approximately 20 million reads were generated per sample.

Reads quality was assessed using FastQC and low-quality sequences removed by using cutadapt with a quality cutoff of 30. The resulting reads were mapped onto the Arabidopsis thaliana genome (TAIR10) using RNA STAR. Mapped reads were counted using featureCounts. Differential gene expression analysis was performed by using the DESeq2 package. A cutoff of p-value adjusted <0.05 and absolute log_2_ fold change > 1 was applied to identity DEGs. RNA-sequencing raw data are available at EMBL-EBI ArrayExpress under the identifier E-MTAB-17058.

### Salicylic acid quantification by UPLC-MS

Leaves of four-week-old Arabidopsis plants were harvested and weighed for fresh weight calculation and immediately flash-freeze in liquid nitrogen. Tissues were ground with a plastic pestle and phytohormones extracted overnight using 0.5-1 ml of ice-cold extraction buffer (methanol: water (80:20 v/v), 0.1% formic acid, 0.1 g/L butylated hydroxytoluene and 100 nM ABA-d6 as an internal standard), as described previously (Roussin-Léveillée *et al*., 2020). Extracted phytohormones were filtered by using several rounds of centrifugation and supernatant collection. Filtered extracts were quantified using an Acquity Ultra Performance Liquid Chromatography system (Waters Corporation, Milford, MA) as described previously. SA was quantified based on a standard curve to calculate sample concentration (nM), which was converted to ng using the molecular weight of each specific compound and the extraction volume used. All data were normalized to initial fresh weight in grams.

### Statistical analysis

Statistical tests were selected as follows. First, normality of the data within each group was assessed using the Shapiro-Wilk test (α = 0.05). If all groups passed normality, homogeneity of variance across groups was evaluated using Levene’s test (α = 0.05). If all groups were normally distributed and variances were homogeneous, a one-way ANOVA was performed followed by Tukey’s Honestly Significant Difference (HSD) post-hoc test for all pairwise comparisons. If all groups were normally distributed but variances were heterogeneous (Levene’s test p < 0.05), Welch’s ANOVA was used instead, followed by Games-Howell post-hoc test, which does not assume equal variances. If one or more groups failed the Shapiro-Wilk normality test, the non-parametric Kruskal-Wallis test was applied regardless of variance homogeneity, followed by Dunn’s post-hoc test with Benjamini-Hochberg correction for all pairwise comparisons. In all cases, a significance threshold of α = 0.05 was applied. Results of pairwise comparisons are displayed on figures as compact letter displays, where groups sharing the same letter are not significantly different. All statistical analyses were performed in R using the base, car, and rstatix packages.

