## Supplementary Figures for "Atypical immunity induced by extracellular water in plants"

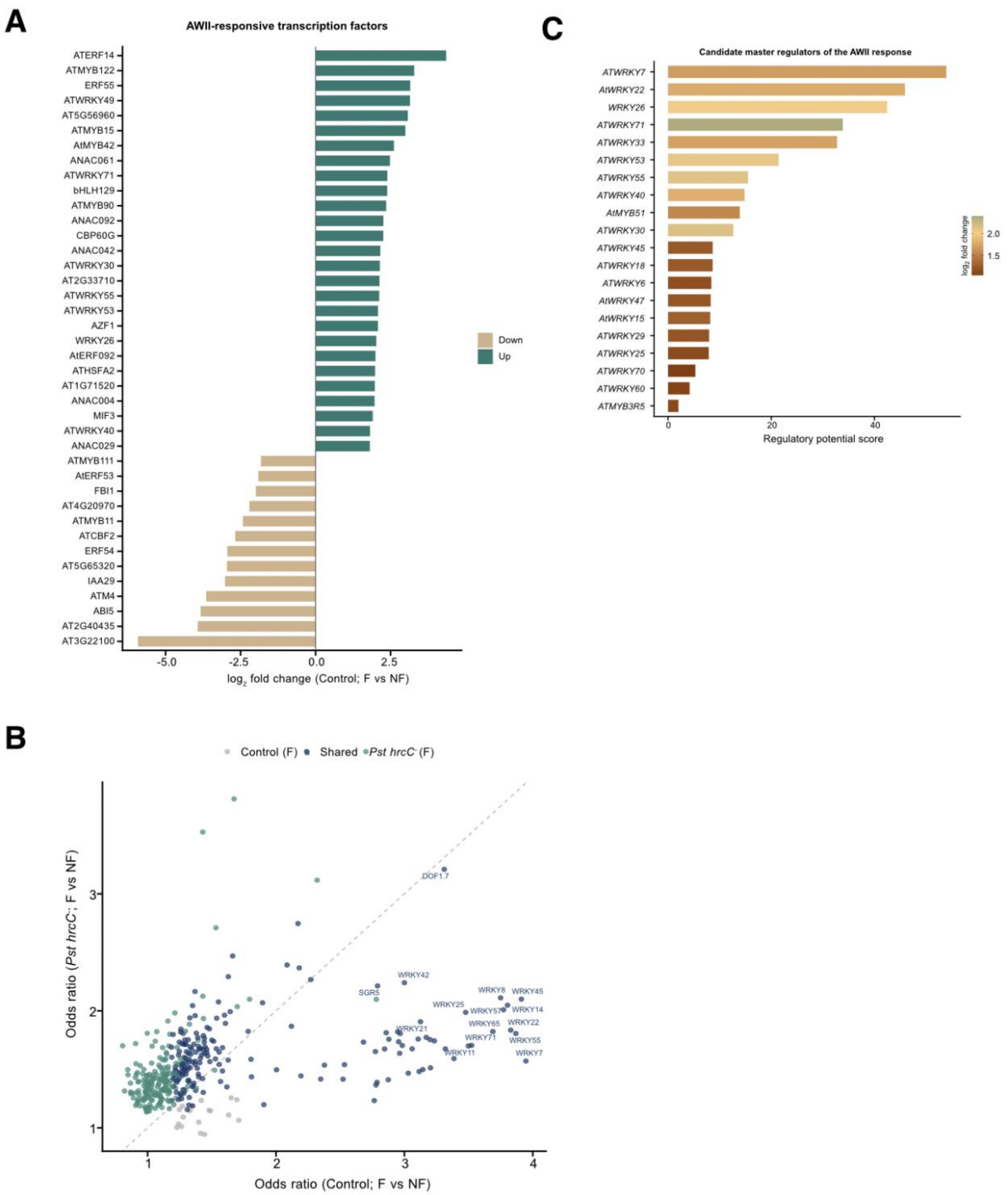


**Figure S1. Gene-regulatory network at the basis of the plant response to extracellular water.** (**A**) Transcription factors differentially regulated in the transcriptome of Arabidopsis WT plants under F state compared to NF state in uninfected (control; $MgCl_2$ 10 mM) plants. Horizontal bar chart showing the top 40 transcription factor (TF) genes ranked by absolute $\log_2$ fold change among DEGs identified in the Control F vs. NF state. In total, 67 TFs were upregulated and 44 downregulated in response to the F state. Bars are colored by direction of regulation: teal (upregulated, $log_2FC > 1$) and brown (downregulated, $log_2FC < -1$). **(B)** Comparison of TF binding motif enrichment between Control and *Pst hrcC*$^-$ F state transcriptomic response. Scatter plot comparing the odds ratios of TF binding motif enrichment in the promoters of upregulated F state DEGs from Control F/NF (x-axis) and *Pst hrcC*$^-$ F/NF (y-axis). Each point represents one JASPAR2022 plant motif (656 motifs tested in total). Points are colored by significance category: shared (enriched in both conditions, padj < 0.05 and OR > 1; n = 177, dark navy), Control-only (n = 18, grey), *Pst hrcC*$^-$ only (n = 188, teal), and not significant in either (n = 273, not shown). The dashed diagonal line indicates equal enrichment in both conditions. The 15 shared motifs with the highest combined odds ratio are labelled with their JASPAR TF name. Enrichment was computed using Fisher's exact test (one-sided) against the background of all expressed genes; p-values were adjusted using the Benjamini–Hochberg. (**C**) Candidate master regulators of the apoplast extracellular water transcriptional response. Horizontal bar chart ranking the top 20 candidate master regulators identified in the Control (F vs NF) state response. Candidates were defined as TF genes that are (1) themselves upregulated (p-value adj < 0.05, $|log_2FC| > 1$) in the Control F vs NF state, and (2) whose cognate binding motif is significantly enriched (padj < 0.05, odds ratio > 1) in the promoters of upregulated flooding DEGs. Candidates were prioritized by a composite regulatory potential score: $log_2FC \times -log_{10}(padj\_motif) \times$ odds ratio.

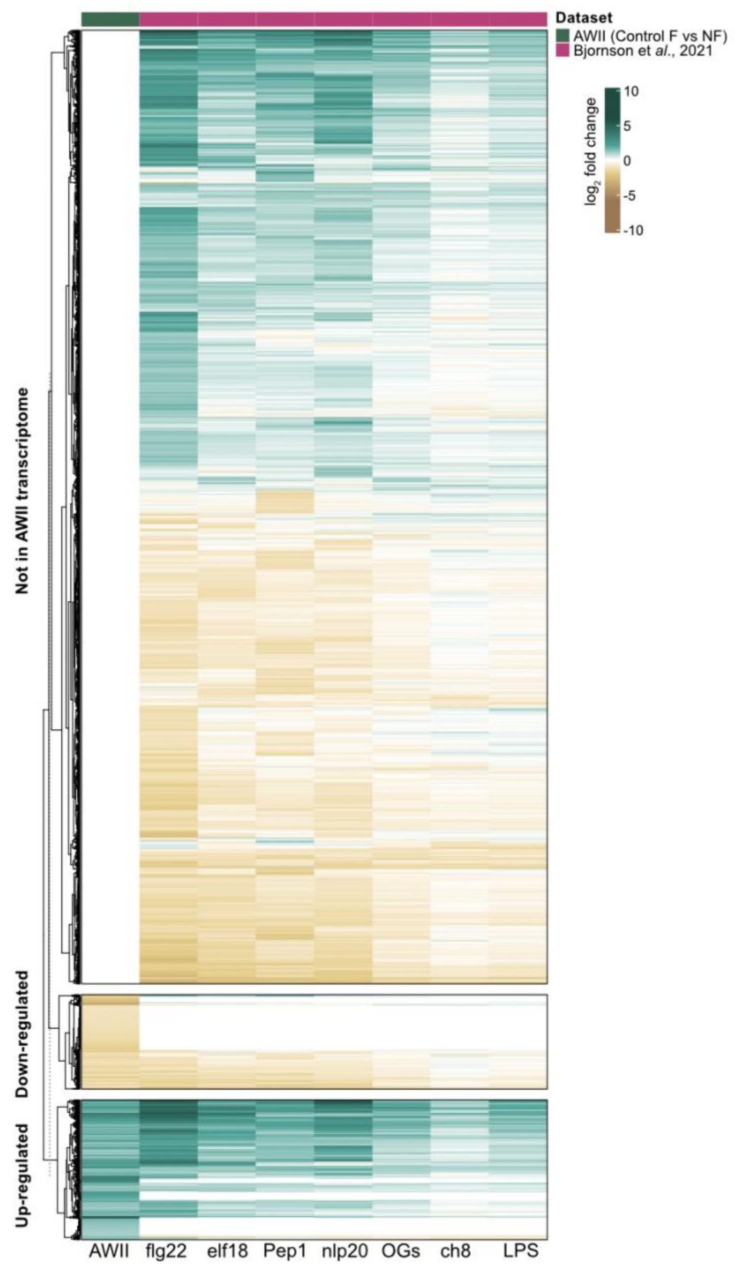


**Figure S2. Comparative analysis between the AWII and PTI/DTI transcriptional signatures.**

Heatmap displaying $\log_2$ fold change values for 8,690 genes representing the full union of DEGs from the AWII transcriptome (control F vs NF state) and the seven PAMP/DAMP-treatment datasets (flg22, elf18, Pep1, nlp20, OGs, ch8, LPS) from Bjornson et *al*. (2021). Rows are divided into three groups: genes upregulated in AWII flooding (1,023 genes, $\log_2FC > 0$), genes downregulated in AWII (691 genes), and genes not present as DEGs in AWII, but significant in at least one PAMP/DAMP-treatment (6,976 genes, "Not in AWII transcriptome"). Row clustering was performed independently within each row group using hierarchical clustering (complete linkage, Euclidean distance).

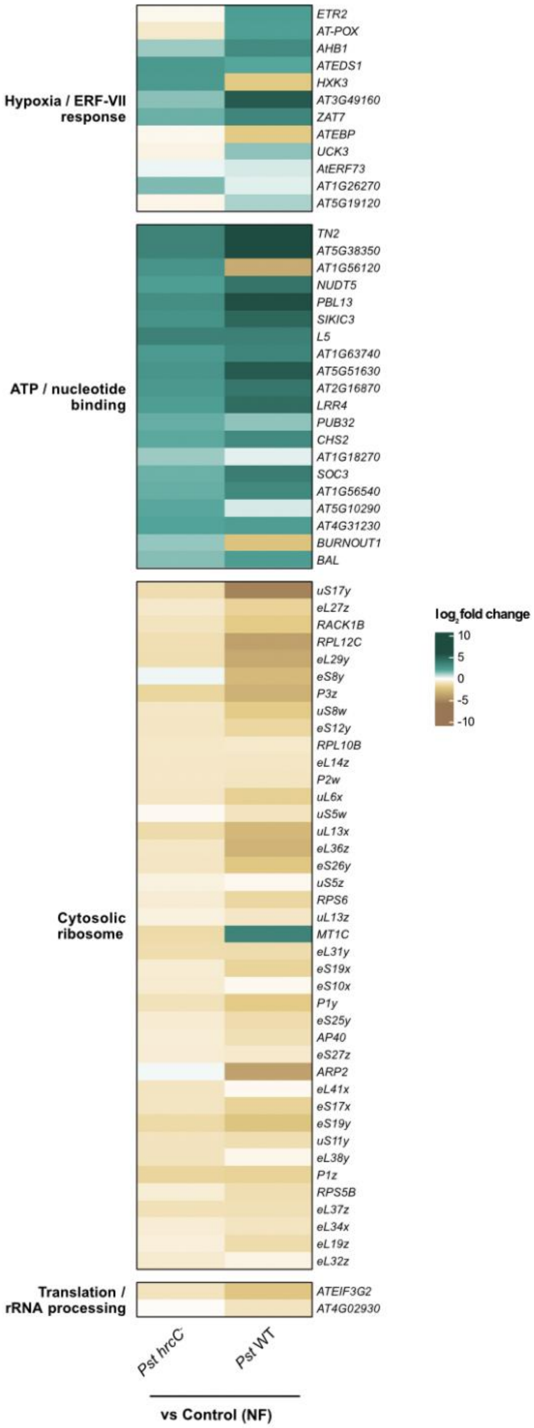


**Figure S3. Non-virulent and virulent *Pseudomonas syringae* infections activate hypoxia-related responses.** Effects of bacteria on the AWII-specific/hypoxia gene sets in NF plants. Heatmap displaying $\log_2$ fold change values for the same 74 AWII-flooding-specific genes shown in Figure 3B, here assessed in non-flooded (NF) conditions. Columns represent two bacterial treatment contrasts within non-flooded plants: *Pst hrcC*$^-$ (NF) or *Pst* WT (NF) vs. Control (NF).

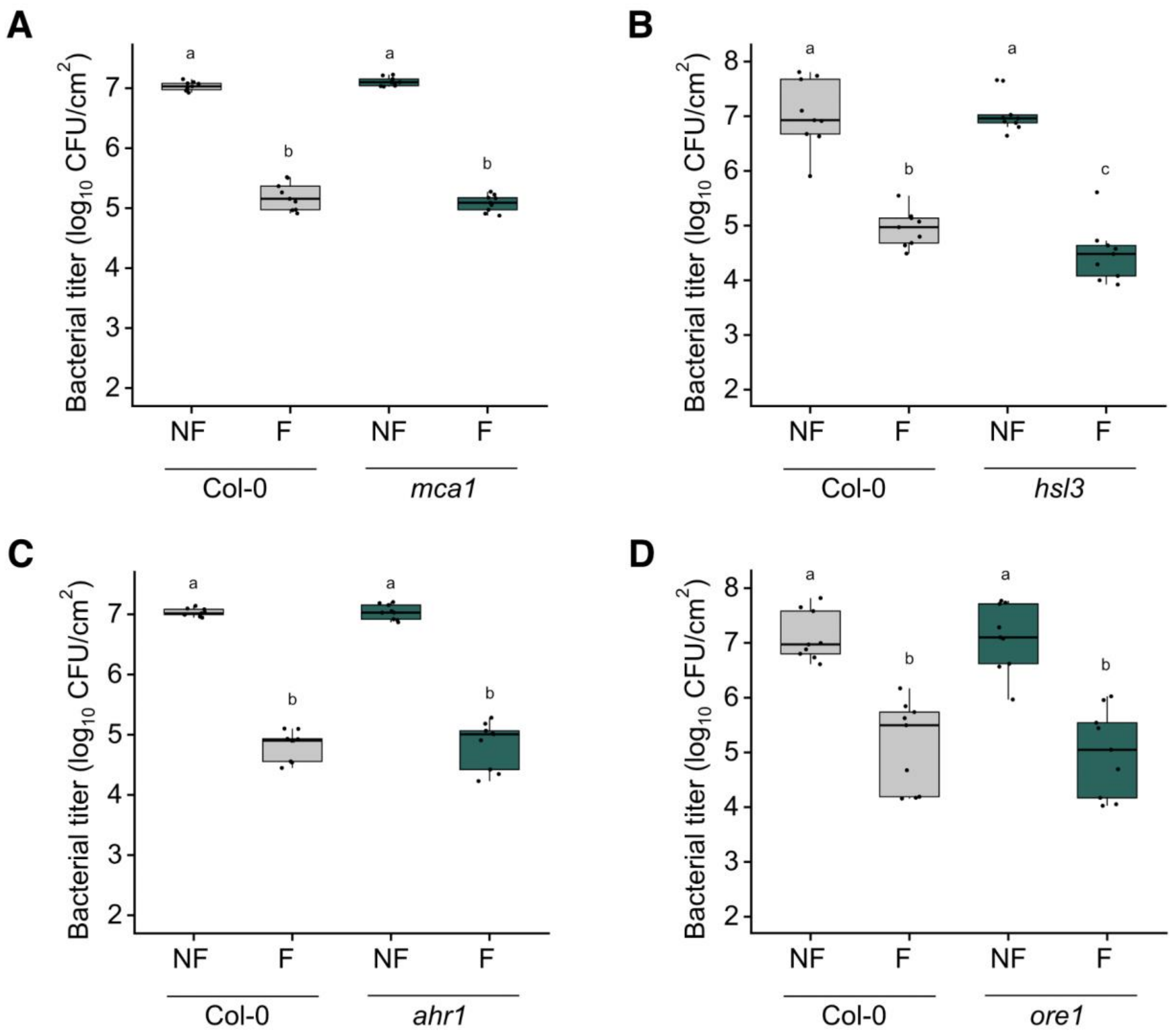


**Figure S4. AWII is likely not triggered by mechano-transduction, SCREW peptide perception, water potential sensing or submergence-induced senescence.** (A-D) AWII is not affected in the Arabdopsis mutants for the (A) *MCA1* (*mca1*; mechano-transduction), (B) *HSL3* (*hsl3*; SCREW peptide perception), (C) *AHR1* (*ahr1*; water potential sensing) or (D) ORE1 (*ore1*; sumergence-induced senescence). Each plant were subjected to AWII (F state for 20 hours) or NF state as performed in 2A, prior to being challenged with *Pst* WT (1 x $10^5$ CFU/ml). Leaves were harvested at 3 dpi for bacterial titer monitoring. Different letters indicate statistical significance ($p<0.05$) obtained after performing a Welch ANOVA followed by a Games-Howell post-hoc test (S4A, S4C), a Kruskal-Wallis test followed by a pairwise Wilcoxon test (BH corrected) (S4B) or a one-way ANOVA followed by a Tukey HSD test (S4D).
